# Activation mechanism of class A GPCRs: machine learning analysis of experimental structural databases

**DOI:** 10.64898/2026.03.26.714415

**Authors:** Santeri Paajanen, Felix Eurasto, Waldemar Kulig, Ksenia Korshunova, Shreyas Kaptan, Ilpo Vattulainen

**Affiliations:** Department of Physics, University of Helsinki, P. O. Box 64, FI-00014 Helsinki, Finland

## Abstract

Recent advances in cryo-electron microscopy and cryo-electron tomography have dramatically increased the number of class A G protein-coupled receptor (GPCR) structures, especially in previously inaccessible G protein-bound, active-like conformations. The increased structural diversity provides a unique opportunity to explore the conformational landscape underlying GPCR activation. To this end, we developed a machine learning (ML) framework that utilizes experimental structural data to elucidate the activation dynamics of class A GPCRs. We find that receptors can populate both inactive and active-like conformations even in the absence of ligand or G protein, providing a structural basis for agonist-free basal activity. Agonist binding shifts this conformational ensemble towards the active state but does not fully stabilize it. Instead, a stable active state is only established upon G protein binding, which locks the receptor in its active conformation. These results support a hybrid activation mechanism in which ligand binding follows conformational selection, while transducer engagement is governed by induced fit. Beyond clarifying class A GPCR activation, the openly available and modifiable ML framework provides a practical tool for analyzing newly determined structures, investigating the mechanisms of action of other GPCR classes and protein families, and guiding structure-based drug discovery in important pharmacological superfamilies.

**Graphical abstract:** 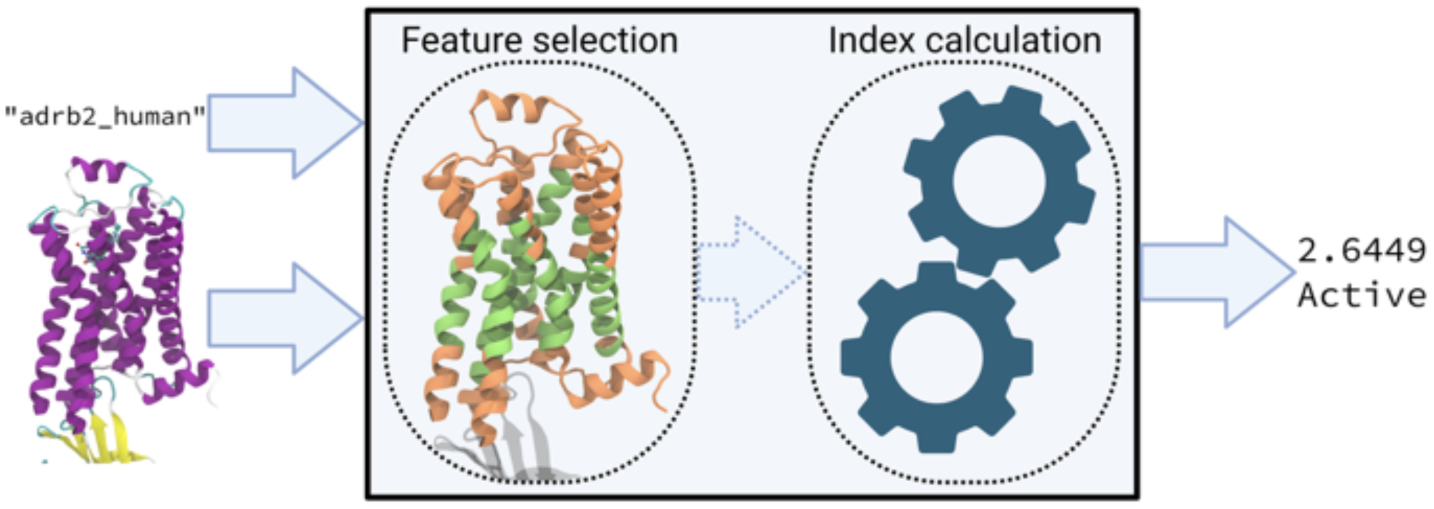

## Introduction

G protein-coupled receptors (GPCRs) are the largest and most diverse family of membrane receptors in eukaryotes. Members of the GPCR family, and especially its subgroup, class A, control essential life functions, including heart rate and numerous brain functions. Given this, it is not surprising that they are a major drug target: about 30-40% of drugs on the market are aimed at controlling their function (Majumdar et al., 2024).

Biomolecular simulations and artificial intelligence (AI) have revolutionized drug development by accelerating target identification and optimizing molecular design (Wei and McCammon, 2024). However, this would not be possible without experimental data of receptor structures.

The structural characterization of class A GPCRs has advanced significantly during the current decade, facilitated by the development of cryo-electron microscopy (Cryo-EM) and cryo-electron tomography (Cryo-ET). This is highlighted in Fig. 1a,d, where we show the results of our analysis of the Protein Data Bank (PDB) on the increasing number of resolved structures of class A GPCRs, highlighting the impressive role of electron microscopy techniques. These techniques have enabled the determination of biomolecular structures close to their physiological state (de Oliveira et al., 2021; Duan et al., 2024), resulting in the deposit of hundreds of active-state structures of class A GPCRs in the PDB (Fig. 1b). Many of these structures include a G protein bound to the receptor (Fig. 1c) and can therefore be used to study the active state of GPCRs.

Activation of GPCRs occurs in a dynamic energy landscape where the receptor conformation does not necessarily change from inactive to active as a result of a single ligand-binding event, but may be a more complex process, proceeding in steps, with ligand binding being only one part of a complex chain of events (Kumar et al., 2000). Although there is a risk of interpreting this complex process in an oversimplified manner, there are currently two main paradigms for GPCR activation. According to the *induced fit* mechanism (Hauser et al., 2021; Koshland, 1958; Latorraca et al., 2017; Manglik & Kruse, 2017; Tehan et al., 2014), GPCR activation is initiated by an extracellular signal, usually in the form of ligand binding to an inactive receptor. Ligand binding then initiates the main structural changes associated with activation, such as the twisting and outward movement of transmembrane helix 6 (TM6, see Supplementary Fig. 1) on the cytosolic side, and the twisting and inward movement of TM5 and TM7. These structural changes induce a change in the signaling protein (usually G protein, arrestin, or GPCR kinase) bound to the cytosolic side of the cleft, which activates signaling cascades inside a cell (Gurevich & Gurevich, 2017; Wang et al., 2018). Recent findings are consistent with the induced-fit mechanism for some receptors, like β2-adrenergic receptor (Asadollahi et al., 2025).

An alternative paradigm is the *conformational selection* mechanism (Ma et al., 1999), in which the protein conformation would spontaneously switch between inactive and active states, even in the absence of ligand, but ligand binding would shift the conformational equilibrium from an inactive to an active state, thereby fostering the stability of the active state.

**Fig. 1.**
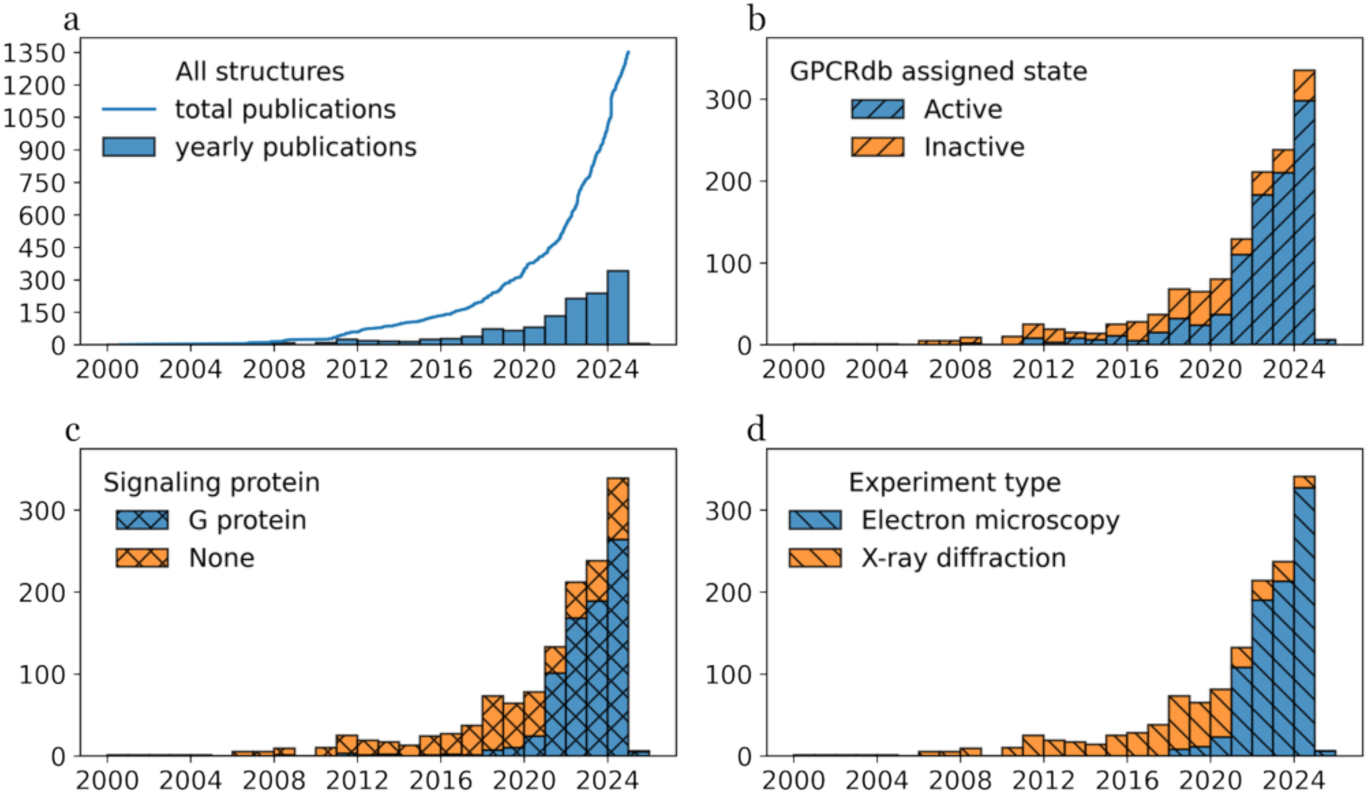
Growth in the number of class A GPCR structures deposited in the PDB. The evolution of the number of cryo-EM structures is reflected in the increase in the number of active structures with a bound G protein, while the evolution of the relative number of inactive structures is decreasing. Data identifiers are from GPCRdb, as listed for each structure. **a** All class A GPCR structures deposited in the PDB: cumulative total number over time (blue line) and deposits per year (bars). **b** Annual deposits broken down by the activation state reported by GPCRdb, considering “active” (in blue) and “inactive” (orange) states. Structures reported as “intermediate” or “other” have been ignored because their number is too low to be distinguished from the graph. **c** Annual deposits broken down by whether the receptor is bound to a signaling protein (G protein, in blue) or not (orange). Arrestins have not been included in the analysis because there is not enough data to influence the graph. **d** Annual deposits broken down by experimental technique, highlighting the importance of electron microscopy (blue) and X-ray diffraction (orange) in structure determination. There were not enough structures produced by electron crystallography to make their contribution concretely visible in the graph.

Although there is more support for the induced fit mechanism (Asadollahi et al., 2025), both paradigms can lead to the same functional state. Regardless of which paradigm is valid, elucidating the dynamic activation processes of GPCRs is in principle already possible using AI-based methods. The potential of AI-based approaches in structure-based drug discovery has already been demonstrated (Michino et al., 2025). However, progress in elucidating the activation mechanisms of GPCRs largely depends on the availability and quality of training data needed for AI analysis. This requires a large number of protein structures that represent different conformational states (inactive, intermediate, and active) and are sufficiently accurate, as well as reliable identification and annotation of these states.

A typical way to determine the activity state of each GPCR conformation is to utilize structural data from experimentally observed active and inactive states, and to complement these experimental data with detailed biomolecular simulation results. If a comprehensive sample of protein structures is available, and if the activity state of each structure has been correctly defined (*e.g*., inactive, intermediate, active), then the activation index obtained from this statistical structural analysis may provide a reliable estimate of the conformational state of each protein. In this spirit, one of the commonly employed models often used as the standard is the so-called A^100^ index (Ibrahim et al., 2019), which is a linear combination of five inter-helix distances developed for class A GPCR proteins. It was trained on simulation data and validated on 268 experimental structures from the PDB, and it has been successfully used in GPCR analyses (Calderon et al., 2023; Alfonso-Prieto and Capelli, 2023). An alternative approach is provided by GPCRdb (Pándy-Szekeres et al., 2018), which calculates the percentage of activation. In this model, structures are classified as fully inactive if a condition based on a certain distance between TM2 and TM6 is met, while fully active structures are G protein or arrestin-bound structures (Kooistra et al., 2021). Otherwise, the percentage of activation is calculated based on the structural similarity to one of these two categories.

However, these and other related models were developed before or shortly after 2020, when there was limited, or in many cases no, structural information on the active state. It is also not certain that the active states of the structures used in developing the models were correctly identified.

In this article, we show that the current structural databases for class A GPCR proteins are comprehensive enough to infer the steps of their activation mechanisms. The machine learning (ML)-based model we developed utilizes only experimental structural data from the PDB, thus avoiding the sampling limitations of atomistic simulations. However, the key strength of the ML model introduced and used here and the main reason for its ability to successfully classify experimentally determined structures is that it utilizes only protein backbone coordinates from conserved structural features of the receptor transmembrane domain, ensuring its general applicability. The model is based on more than 1000 GPCR structures and trained in an unsupervised manner, providing a robust and rigorous technique for analyzing experimental structural databases.

The results confirm that ligand binding is the first event in a stimulated activation sequence. Furthermore, the results reveal conformational changes that are common among these receptors—inward movement of TM7 and outward movement of TM6—that occur during activation and precede the binding of G protein to the receptor. We find that most of the experimental structural data was originally interpreted correctly in terms of states corresponding to the observed structures, but the ML model also identifies anomalous structures whose state of activity was originally interpreted incorrectly. This leads to a key observation, the *apo* form of several class A GPCRs exists in both the inactive and active states, and the agonist-bound form also exists in both the inactive and active states. This indicates that the conformation of class A GPCRs spontaneously switches between the inactive and active states in both the absence and presence of ligand. Ligand binding stabilizes the active state, but G protein is required to lock the receptor into the active state conformation. Experimental structure data hence reveals that class A GPCR activation is mainly regulated by conformational selection at the ligand level and complemented by the induced fit mechanism at the transducer level. The ML model used in the analysis is openly available as a tool that can be used to analyze new experimental structures and test the validity of predictions made by foundation models, such as AlphaFold.

## Results

### The index G_CA_ distinguishes between active and inactive structural classes

Using principal component analysis (PCA), we developed a new structural index G_CA_ (see Methods) that reveals the different structural classes underlying the class A GPCR structures deposited in the PDB. Based on this index, Fig. 2 presents the main results of this article.

Figure 2 shows that the G_CA_ index distinguishes antagonist-bound structures from structures bound to both G protein and agonist (orange line in Fig. 2a vs. blue line in Fig. 2b and orange dots *vs.* blue dots in Supplementary Fig. 2a). This confirms that the index serves as a reliable proxy of receptor activation. The distribution of structures described by the index allows for classification into two functional states (see especially Supplementary Fig. 2a). Structures bound to both G protein and agonist (Fig. 2b, blue line) are predominantly found with index values above –1.72, while G protein-free and antagonist-bound structures (Fig. 2a, orange) are mainly found below this threshold. Thus, index values above – 1.72 define the active region, while values below this threshold define the inactive region.

**Fig. 2.**
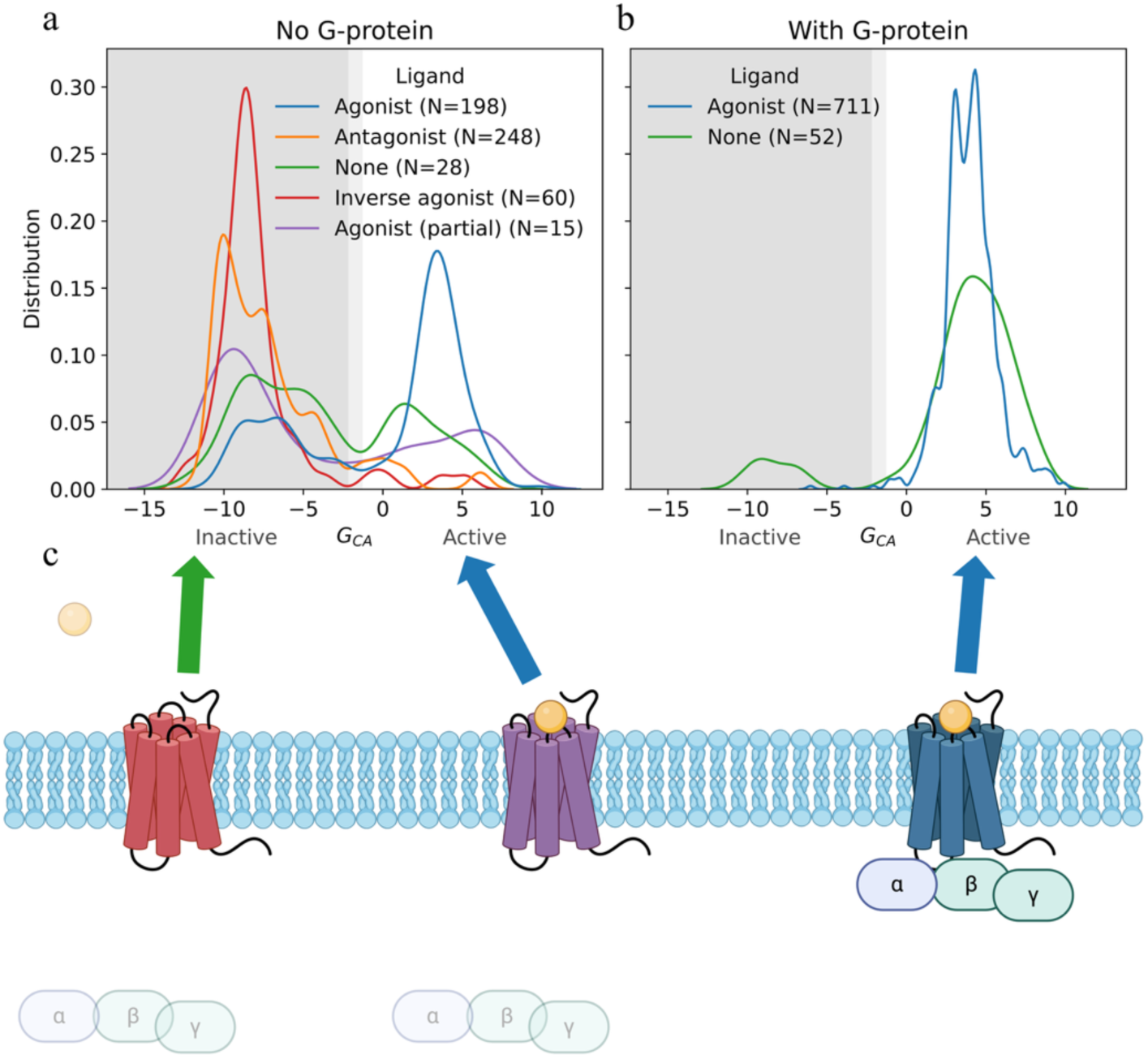
Distribution of class A GPCR structures in the PDB in the inactive and active states, as described by the index G_CA_. **a** Distribution of class A GPCR structures as indicated by the index G_CA_, showing the distributions for different ligands in structures without G protein binding. The inactive region is shown in gray. The light gray region depicts the 95% confidence interval. *N* is the number of structures analyzed. **b** Effect of ligand on the distribution as indicated by the index in structures with G protein binding. Inactive and active regions are labeled as in panel a. **c** Schematic mechanism of GPCR activation illustrated from left to right: inactive GPCR before ligand and G protein binding (left), ligand-bound activating state (middle), and ligand and G protein bound stable active-state conformation (right).

In this framework, structures bound to both G protein and agonist simultaneously correspond to the active state of the GPCR. Structures bound only to antagonist (without G protein) are predominantly in the inactive state. Interestingly, structures bound to G protein but not to agonist, and structures bound to agonist but without G protein, are distributed over a broad front along the index, although both have a peak in the active region that represents the primary state. In contrast, G protein-bound or ligand-bound structures do not have a dominant state in the inactive region.

The results show that the index G_CA_ reliably describes the activation state of class A GPCRs and can be used as a quantitative descriptor of functional conformations.

### Activation is driven by the cytosolic inward movement of TM7 together with the outward movement of TM6

The outward movement of TM6 is the most important indicator of GPCR activation (Hauser et al., 2021; Latorraca et al., 2017; Manglik & Kruse, 2017). This feature is a common denominator in structural data and is also reflected in our index. Using the ML model and interpolating between the structures it describes, from inactive to active states, we find that the outward movement of TM6 is coupled to the inward movement of the cytosolic terminal of TM7 (see Supplementary Movie). The movement of TM7 is almost perpendicular to the outward movement of TM6. On the cytosolic side, there are also smaller outward movements of TM1 and TM2 and inward movements of TM3 and TM5, while TM4 rotates only slightly in place. These smaller changes of TM1, TM2, and TM4 during larger movements of TM6 and TM7 may be due in part to the removal of rotation and translation required by the ML analysis. It is worth noting that this does not affect the calculated index values, but only the interpretation of the interpolated conformational changes (Supplementary Movie), as the additional rotation becomes a part of the model. On the extracellular side, no significant differences in the index between the two states are observed, only a small shift in TM3, indicating that the detailed activation mechanisms of the ligand binding sites are not shared among the different receptors.

### Agonist binding alone is sufficient to drive the receptor to the active state, but G protein is needed to stabilize this state

The distribution of experimental structures reveals that receptors bound to an agonist in the absence of G protein preferentially occupy the active region (Fig. 2a). This supports the top-down view of activation, in which agonist binding to a receptor promotes activation and promotes the stability of conformations associated with receptor activation. While this behavior is consistent with the conformational selection mechanism, it may also be compatible with the induced fit model, as the structural distributions do not reveal whether the ligand initially binds to the inactive or active receptor conformation. Yet the observation that *apo* structures are also found in the active region (Fig. 2a) suggests that class A GPCRs can spontaneously adopt active-like conformations, and they can switch between inactive and active-like conformations, even in the absence of ligand. This is in line with the conformational selection mechanism rather than the induced fit model and is consistent with the basal activity observed for many GPCRs (Seifert et al., 2002; Tao et al., 2008; Hammes et al., 2009; Vogt et al., 2012).

The role of antagonists, however, is more nuanced: rather than simply blocking the effects of an agonist or inverse agonist, antagonists attenuate the dynamics of their receptor along the activation coordinate, stabilizing the inactive state. This contrasts with the common definition of antagonist, which simply interprets antagonists as merely blocking the effects of agonists and inverse agonists, and emphasizes the active role of antagonists in modulating receptors’ conformational landscapes.

This view is supported by the fact that the distributions of antagonist-bound and inverse agonist-bound structures (neither bound to G protein) differ significantly (*p* value of 0.018 in a two-sample Kolmogorov-Smirnov test). The results (Fig. 2a) suggest that inverse agonists act as stronger inactivators than antagonists, blocking the receptor and locking it more effectively in the inactive state. In contrast, partial agonist-bound structures show a lower tendency to occupy the active region compared to agonist-bound structures, suggesting their inefficiency, consistent with their biochemical classification.

### Index anomalies point to mislabelled structures

The structural classifications provided by the index revealed a small subset of G protein-bound structures (8HIX, 8HJ0, 8HJ1, 8HJ2, 8HMP, 7XBW, 7XBX) that are located in the inactive region of the activated space (Fig. 2b). All these structures are marked as active in the GPCRdb classification. However, a closer look at the G protein-binding site shown in Fig. 3 reveals that these structures share key features of the inactive conformation: TM6 remains close to the center of the helical bundle of a GPCR, while TM7 is shifted outward—both being hallmarks of the inactive state. Since TM6 physically blocks the G protein-binding pocket in this conformation, these structures appear to be based on structural anomalies that still allow G protein binding. The gpr21 structures 8HIX, 8HJ0, 8HJ1, and 8HJ2 (Lin et al., 2023) are missing a segment of TM6 that would normally clash with G protein (Fig. 3b). In contrast, in the gpr52 structure 8HMP (Fan et al., 2023) (Fig. 3c) this part is present, but it is kinked, suggesting that G protein is adapted for binding through structural distortion, possibly through forced insertion. Similarly, in the cx3c1 structures 7XBW (Fig. 3d) and 7XBX (Lu et al., 2022) the receptor itself is in the canonical inactive conformation, but the G protein helix has rotated clockwise, and its C-terminal fold has partially unfolded.

These structural anomalies suggest that there is no reason to expect G protein binding to receptors in an inactive conformation. Such interactions, if observed, are likely to be due to experimental artifacts, construct design, or nonphysiological stabilization methods. This supports the view that receptor activation—and the associated structural rearrangement, particularly the outward movement of TM6—is a prerequisite for G protein binding.

### Comparison to previous PCA-based analysis of GPCR structural data

A previous study (Wingert et al., 2022) performed PCA on 160 class A GPCR structures. The analysis revealed similar subclasses, especially among human receptors. However, they also reported a clear species-dependent separation into different subclasses, suggesting possible differences in activation mechanisms or structural configurations between species. Due to the significant expansion of the dataset in 2022-2025 (see Fig. 1 and Methods), our analysis no longer indicates this species-specific distinction (Supplementary Fig. 2). Instead, we find that bovine receptors, which are exclusively sensory receptors in the dataset, cluster with a small number of available human sensory receptors. This finding suggests that the receptor subgroup has a stronger influence on structural clustering than the species of origin. Furthermore, in our analysis, sensory receptors show a clearer distinction between active and inactive conformations and share a common distribution and G_CA_ index threshold. This consistency supports the reliability of the G_CA_ index developed in this work to describe the activation state of class A members, regardless of species or receptor subclass.

It is worth emphasizing the importance of using comprehensive datasets in developing generalizable and diverse GPCR activation models. The requirement for diversity encompasses both functional coverages, with samples of active and inactive structures, and different receptor types, with structures from different subtypes and species.

## Discussion and conclusions

The basis for understanding the function of molecular motors and receptors is knowledge of their structure and its changes during activation. For membrane proteins, this challenge has been problematic, as elucidating their structures has been painfully difficult. New techniques based on cryo-EM and cryo-ET have been a game changer in this regard. Combined with the use of nanodiscs, they produce an increasing number of membrane protein structures, which has made protein structure databases a rich source of information for a more comprehensive understanding of membrane protein function. However, the information they contain is surprisingly underutilized.

In this work, we developed a machine learning-based model to analyze the structural data for class A GPCRs in the PDB. The goal was to investigate whether the amount of information currently available in the databases can already be used to elucidate mechanistically how proteins in this family are activated.

**Fig. 3.**
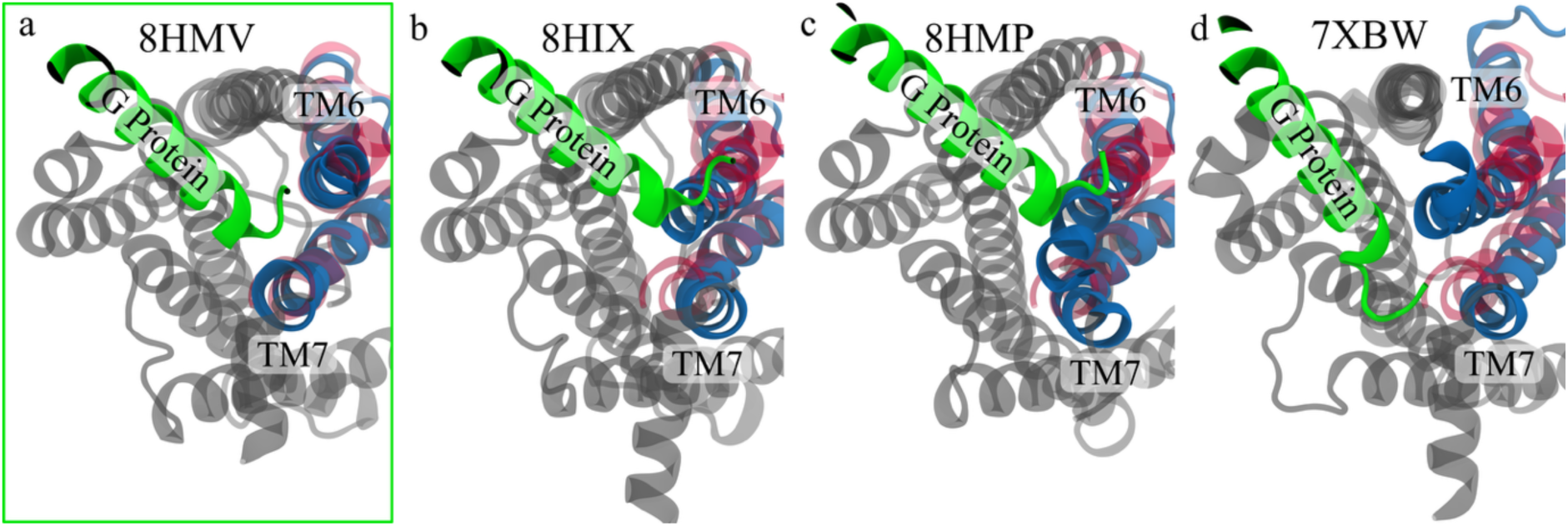
Visualization of experimentally determined protein structures, highlighting the differences between the aberrant inactive structures and the genuine active structures. The C-terminal helix of the G protein alpha subunit is shown in green, TM6 and TM7 of the experimentally determined (but misinterpreted in terms of activity state) protein structures in blue, and TM6 and TM7 of the genuine active-state structure (3P0G) in red. **a** Genuine active structure of gpr21 (8HMV). **b** Inactive structure of gpr21 (8HIX). TM6 is turned inward as in the inactive structure, which should intersect with the G protein helix, but the end is not included in the structure. The same defect is evident in 8HJ0, 8HJ1, and 8HJ2. **c** Deformed inactive structure of gpr52 (8HMP). The G protein helix pushes on TM6, bending it inward and towards TM7. **d** Inactive structure of cx3c1 (7XBW). The G protein helix is rotated clockwise to avoid a clash with TM6, and the end of the G protein helix is partially unfolded to get closer to TM7. The same defect is evident in 7XBX.

The analysis confirms that there are two distinct states for the inactive and active conformations. The structural difference observed between these two states is consistent with the previously observed GPCR activation mechanism (Zhou et al., 2019; Latorraca et al., 2016), which is characterized by the movement of TM6 as it swings out of the helical bundle and TM7 swinging in during activation. However, detailed analysis of the structural data reveals that the events that drive class A GPCRs into the active and inactive states are less obvious, and ML analysis provides insight into the activation mechanism of these receptors.

In cryo-based microscopy experiments, the protein structure to be determined corresponds to an ensemble average of numerous structures. In this case, it is often assumed that in the case of an *apo* form, this structure corresponds to the inactive state, while in the case of an agonist-bound state, the active state would predominate. Contrary to this expectation, however, we found that the *apo* form of several class A GPCRs exists in both the inactive and active states. Similarly, we found that the agonist-bound form also exists in both the inactive and active states. The distributions of the *apo* and agonist-bound forms are bimodal, with the dominant peaks in the inactive and active states, respectively. However, in both cases, the observed minority peak is significant, indicating that the conformation of class A GPCRs spontaneously switches between the inactive and active states in both the absence and presence of ligand.

These findings shed light on a controversial question regarding the mechanism of activation of class A GPCRs, namely whether activation occurs through the induced fit mechanism or the conformational selection mechanism (Hammes et al., 2009; Vogt et al., 2012). According to the induced fit hypothesis, the driving force is ligand binding to the receptor, which results in a conformational change of the protein from an inactive conformation to one corresponding to the active state. The conformational selection mechanism, on the other hand, assumes that the target protein undergoes states according to an equilibrium distribution of inactive and active states, and ligand binding shifts the equilibrium to favor the active state.

Two important features are clearly revealed by the structural data analyzed with our ML model. First, on the one hand, a ligand-free protein can reach a conformation corresponding to the active state, and on the other hand, a ligand-bound protein can be in a conformation matching the inactive state. Second, when we studied the structures of G protein-bound receptors, we found that they almost unambiguously represent only the active state, suggesting that the function of G protein is to lock the receptor in the active state.

Thus, experimental structural data on class A GPCRs suggest that, with respect to the ligand, these GPCRs follow the conformational selection mechanism, and with respect to G protein, class A receptors follow the induced fit mechanism.

These findings have broad implications for the pharmacology and drug development of GPCRs. First, the existence of (albeit few) active conformations in the *apo* state provides a structural basis for the constitutive (basal) activity commonly observed in class A GPCRs (Seifert et al., 2002; Tao et al., 2008). This inherent conformational plasticity explains why some receptors exhibit significant signaling in the absence of agonists and highlights how disease-associated mutations can increase basal activity by altering the balance between inactive and active states.

Second, conformational selection for agonist binding suggests that ligands may promote the stability of conformations corresponding to the active state, rather than inducing new conformations starting from a purely inactive state. This mechanism supports the rational design of agonists that exploit the natural energy landscape of the receptor (Deupi et al., 2010), potentially improving potency and effectiveness by targeting the *apo* ensemble to a smaller active population. It also provides a framework for explaining partial agonism: partial agonists may promote active states less efficiently than full agonists, leading to changes in the conformational distribution during the intermediate phase of activation.

Third, the ability of agonist-bound receptors to sample inactive states highlights the dynamic nature of ligand efficacy and suggests that ligands can modulate the conformational equilibrium between inactive and active states in different ways. This conformational selection model is consistent with the molecular basis of biased agonism (functional selectivity), in which structurally different ligands stabilize overlapping but non-identical conformational distributions (Thomas et al., 2026) and preferentially couple to G proteins, β-arrestins, or other transducers. Our findings thus provide structural evidence that biased signaling is due to ligand-specific population changes in the activation landscape, rather than to completely distinct ligand-induced conformations.

Finally, the dual mechanism observed in this work, *i.e.* conformational selection of orthosteric agonists and induced fit for G protein binding, has direct implications for therapeutic design. Agonists that are strongly biased towards active states may facilitate more nuanced G protein coupling while allowing for physiological regulation. On the other hand, the role of G proteins in locking the receptor into its active state conformation suggests that strategies to stabilize the active states bound to the transducer (*e.g*., using nanobodies, conformation-stabilizing antibodies, or bitopic/allosteric modulators) could improve the efficacy or lifetime of signaling. This perspective supports new approaches in GPCR drug discovery, such as conformer-selective screening, allosteric modulation of activation equilibria, and the development of biased ligands to achieve pathway-specific effects and reduce on-target side effects.

In summary, the analysis of experimental protein structures performed in this work reveals that class A GPCR activation is mainly regulated by conformational selection at the ligand level, complemented by the induced fit mechanism at the transducer level. These findings refine models of receptor dynamics and provide new opportunities for the structure-guided design of more selective, efficacious, and safer therapeutics targeting this important pharmacological class. The Open Access tool developed here is available and modifiable to studies of other GPCR classes and other protein families, with extending structure databases, providing a valuable resource for the structural biology community.

## Methods

The methodology is described below in a way that highlights the main points. Certain details will be added once the article has been accepted for publication.

### Data collection and featurization

The structural data used in this work were collected from the curated GPCRdb database (Isberg et al., 2015). We collected all class A GPCRs from GPCRdb, assembling a total of 1010 PDB structures, filtering out only those that did not have a downloadable PDB file in the RCSB PDB.

To featurize the data, we used the generic numbering of GPCR residues provided by GPCRdb. We first retrieved a list of published protein structures and their PDB IDs from GPCRdb, and then, using the Uniprot protein ID included in the answer, we re-retrieved the generic numbering of each protein from GPCRdb. Finally, the structure PDB file was downloaded from the RCSB PDB (Berman et al., 2000) and a pairwise alignment of the structure and protein sequence was performed using the *PairwiseAligner* tool in *Biopython* (Cock et al., 2009). The parameters used were a match score of 2, a mismatch score of −1, and an internal_gap_open_score of −10. All other options were left at the default value of −1. These values represent the added score for each matching amino acid residue, a penalty for mismatches, and a penalty for gaps, respectively. Only contiguous parts of the structures’ sequences (according to residue numbering, ignoring gaps less than 10 in length) with at least 10 amino acid residues were independently aligned to the protein sequence. If the matched score was less than half the sequence length, the part was considered an addition to the sequence and was ignored.

Once the structures were aligned, generic numbers that appeared in all structures were selected for further analysis. A schematic representation of the alignment and number selection is shown in Fig. 4a. This procedure produced a set of amino acid residues that broadly covered all seven transmembrane helices and all class A GPCRs. Figure 4b illustrates the selected residues, highlighting the good coverage especially in the transmembrane region.

**Fig. 4.**
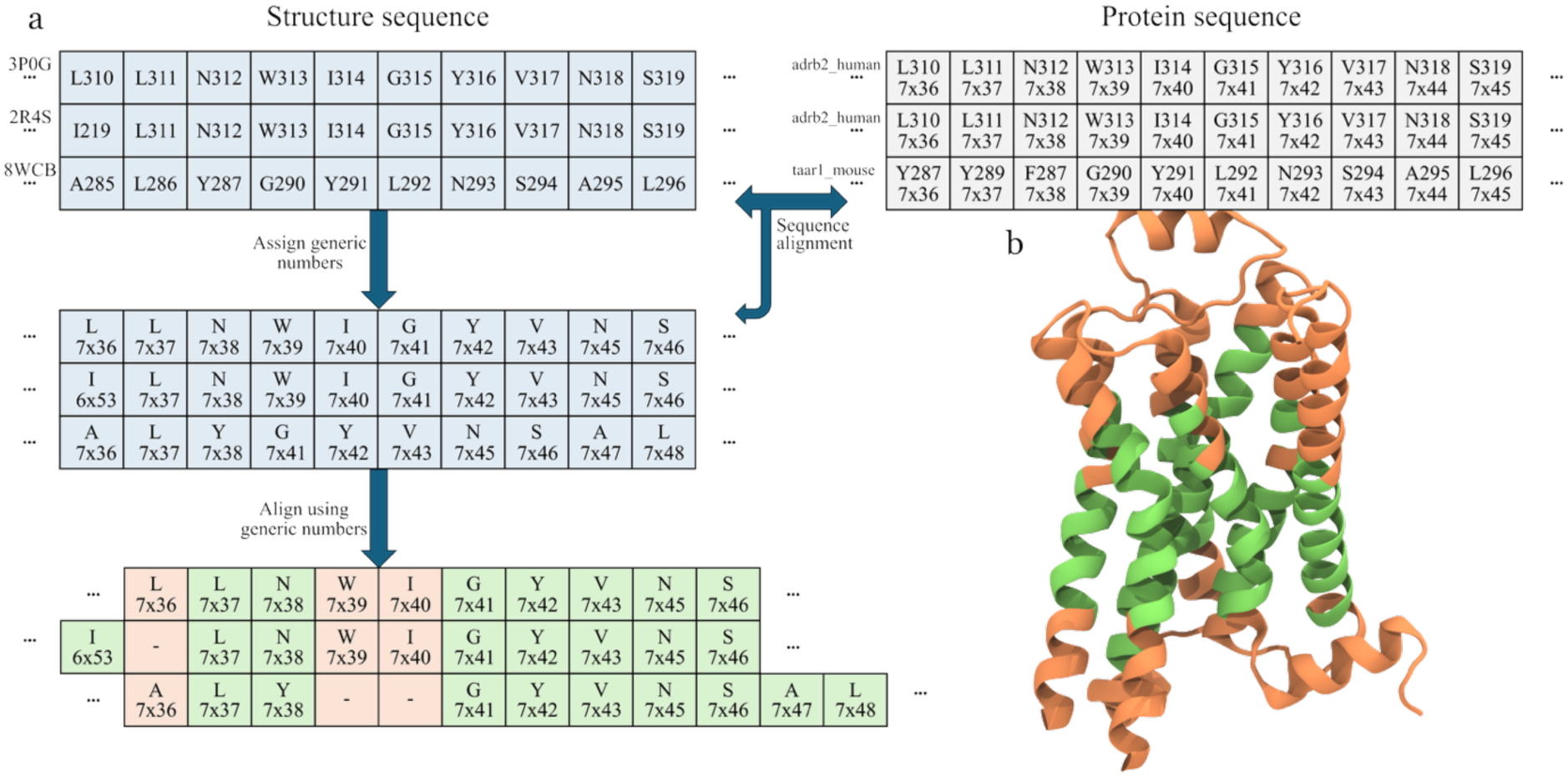
Structural model. **a** Schematic diagram showing the structure alignment process using the 3P0G, 2R4S, and 8WCB sequences as examples. First, the sequence of each structure was assigned a generic number by aligning the sequence to a protein sequence with known generic numbers. These generic numbers were then used to align the different sequences of all structures. Only generic numbers found in all structures were used for further analysis. **b** PDB structure of 3P0G showing which parts of the transmembrane helices were selected for the analysis. Green regions had corresponding amino acid residues in all structures, while orange regions were missing at least one or had no generic numbering. The analysis was done on the alpha carbon coordinates of those residues that are not missing in any structure.

### Dimensionality reduction and interpretation

Using the coordinates of the selected alpha carbons, we applied PCA to reduce the dimensionality of the data. Rotations and translations were removed using the SVDSuperimposer function of the *Biopython* package (Cock et al., 2009) using the structure 3P0G (Rasmussen et al., 2011) as the reference. Coordinates were written and read in the XTC file format (Abraham et al., 2015) using the *MDTraj* package (McGibbon et al., 2015). PCA was calculated using the *scikit-learn* package (Pedregosa et al., 2011). The contribution per component after the first five components (*i.e*., how much the direction of each component explains the total variance in the data) falls below 5%, indicating that the amount of information in the principal components after the first five components is of little importance (Supplementary Fig. 3). Supplementary Fig. 4 shows the projections between the first five PCA components, and from it we can see that only the first component distinguishes antagonist-bound *vs.* G protein and agonist-bound structures. This strongly suggests that only the first component indicates activation.

After training the model, the GPCRdb query was rerun, adding new structures published during the training phase to the analysis and updating some of the labels. The selection was kept static, and the model was not retrained with new data. Eight of the new structures were ignored because they were missing a residue from the list. All receptor classes presented in this article refer to this version (Supplementary Table 1–3), and the number of structures analyzed in this work also corresponds to these selections (Supplementary Table 1–3). Supplementary Fig. 5 summarizes the analysis of the new structures. The distributions in Fig. 2 were calculated as kernel density estimates using the kdeplot function of the *seaborn* package (Waskom, 2021), with the bw_adjust parameter set to 0.5 to achieve a higher resolution distribution. The underlying histograms are presented in Supplementary Fig. 6. The difference between the agonist- and antagonist-bound structure distributions was quantified with a two-sample Kolmogorov–Smirnov test using the stats.ks_2samp function of the *scipy* package (Virtanen et al., 2020).

### Structural index G_CA_

We define a novel activation index (G_CA_; referring to the context ‘<u>G</u>PCR <u>C</u>lass <u>A</u>’) that exploits PCA using only empirical structural data. To calculate the index, we use the coordinates of the alpha carbon atoms of the selected amino acid residues. The 1010 analyzed structures were aligned both rotationally and translationally to a reference structure (3P0G) (Rasmussen et al., 2011) using the SVDSuperimposer function of the *Biopython* package (Cock et al., 2009). This step minimizes the root mean square deviation (RMSD) between all structures and the reference. The coordinates of the thus aligned structures form a matrix, which was used to build a PCA model using the *scikit-learn* package (Pedregosa et al., 2011). With this information, the index can be calculated for any target structure as follows:

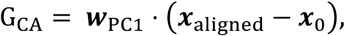

where 𝒘_PC1_ are the weights of the first component of the PCA model, *x*_aligned_ are the aligned coordinates of the target structure, and *x_0_,* are the mean coordinates of all structures, which are calculated as follows:

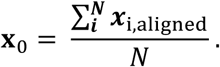

The choice of the first component to calculate the index is based on our previous observation in Supplementary Fig. 4, which shows that the first component is able to distinguish active GPCR structures bound to G protein from inactive ones.

The index can also be determined when some residues in the selected set are missing. A mathematical formulation is given in Supplementary Data. The reliability of the index remains high even then: the error estimate depends on which residue is missing, but even if several residues are missing, the error remains relatively small.

### Clustering

Gaussian mixture models (GMM) (Reynolds, 2009) were used to find explicit categories in the reduced dimensional space. Specifically, we used a one-dimensional GMM on the first principal component and decomposed the data into two classes. To estimate uncertainty of the threshold index, we performed bootstrapping analysis by generating resampled data sets from the original data and rebuilding the GMM. To obtain a robust estimate and minimize the statistical error, we used 10,000 resampled data sets. With a 95% confidence interval, this resulted in a threshold value of –1.72 ± 0.44, with the active region defined as index values greater than this threshold.

### Key strengths of the machine learning model

The machine learning model developed in this work has several significant strengths. First, the model does not include structural details of proteins, such as side chain orientations. This makes it insensitive to the interpretation problems of noisy experimental structural data, which are inevitable if the resolution of the experimental data is not on the order of ∼1 Å throughout. Instead, the model utilizes only backbone information from the structural data about conserved structural features in the transmembrane domain region of the receptors, which is essential specifically for members of class A GPCRs.

Second, the model is based on an established methodology (PCA), which makes its operation robust. The chosen model is a single-component PCA model, combined with a GMM, which makes it easy to interpret. This approach lays the foundation for a general method for estimating the activation level from protein structures, without requiring high resolution or detailed information from the protein structures. The chosen methodology thus works seamlessly with the protein structure mapping method stated above.

The chosen methodology can also be applied to structures that lack a significant number of amino acids, if the alpha carbon positions of the selected specific amino acids of the transmembrane helices required for the calculation are present.

Additionally, given that the model utilizes only experimental structural data, potential problems that might arise from including simulation data as part of the analysis are avoided.

The result is a model that effectively distinguishes between active and inactive states of class A GPCRs using only experimental data. If any experimental structure was initially misinterpreted and labeled, the model can identify and correct it.

## Acknowledgements

We thank the Ministry of Education and Culture of Finland for financial support as part of the doctoral education pilot project (F.E.). We also thank the Helsinki Institute of Life Science (HiLIFE) Fellow program, Sigrid Jusélius Foundation, Lundbeck Foundation, and Academy of Finland (project IDs: 335527, 331349, 336234, 364185) for financial support (I.V.). The authors acknowledge CSC–IT Center for Science Ltd (Espoo, Finland) for providing computational resources.

## Author contributions

S.K. and I.V. conceived the project. S.P., F.E., and S.K. developed the machine learning model. S.P. and F.E. analyzed the data. K.K., W.K., S.K., and I.V. supervised the research. I.V. acquired resources for the project. S.P. and S.K. wrote the first version of the manuscript, and all authors edited the manuscript.

## Supplementary Data

- Mathematical formulation for the determination of the G_CA_ index with missing residues.

## Supplementary Information

- Additional results: Supplementary Figure 1-6, 1-8, Supplementary Table 1-3.

## Supplementary Movie

- **Supplementary Movie: Interpolation of the first principal component (G_CA_ index) from low (inactive) to high (active) values.** The interpolated C-alpha coordinates are shown in a surface representation colored by the magnitude of the component (which is directly proportional to the magnitude of the motion) from white to cyan to magenta to red. Left: shown here are the interpolated coordinates overlayed on the reference structure (3P0G) as a cartoon representation colored by secondary structure (alpha helix in magenta, turn in cyan, and coil in white). Right: a view of the interpolated coordinates from the extracellular side. On the bottom right, there is a view of the interpolated coordinates from the cytosolic side. The largest motions can be observed on the cytosolic ends of TM6 and TM7, with TM6 moving outwards and TM7 inwards.

## Supplementary Data

### Determination of the G_CA_ index with missing residues

The index is calculated from the component 𝒘_PC1_ = (𝑤_1_, 𝑤_2_, …, 𝑤*_n_*) and the aligned and centred coordinates 𝒙 = 𝒙_aligned_ – 𝒙_0_ = (𝑥_1_, 𝑥_2_, … , 𝑥*_n_*) in the form

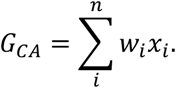

Let the set of missing features be *M*, and the set of non-missing features be *N*. Then the part of the index calculated without missing features is

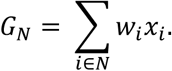

Since we do not know the values of 𝑥_i_ when 𝑖 ∈ 𝑀, we consider it to be proportional to the (estimated) value of the index, 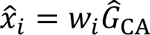, yielding an estimate

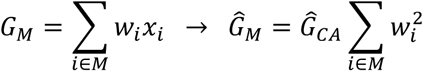

According to the definition of PCA, 𝒘_PC1_ is a unit vector, i.e. 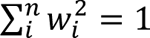. Thus, we get

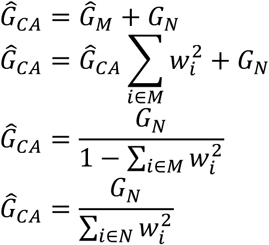

## Supplementary Information

**Supplementary Fig. 1.**
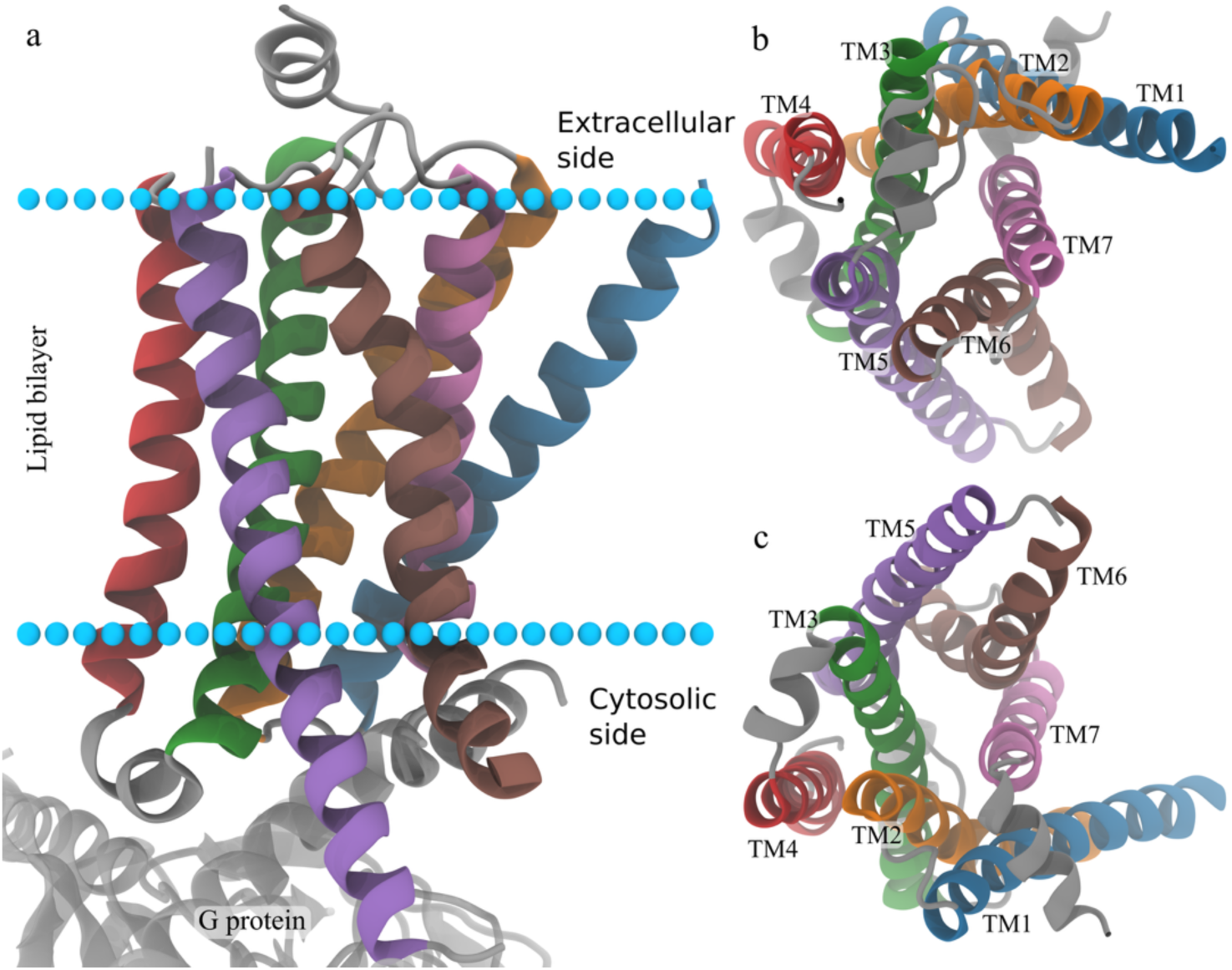
Example structure 3SN6 with labels showing the transmembrane region of the receptor. Transmembrane helices are colored as blue, orange, green, red, violet, brown, and pink for TM1-7, in respective order. Extra- and intracellular regions are depicted in grey. **a** Side view of the receptor with cyan spheres showing the approximate location of the membrane as calculated by the OPM server (Lomize et al., *Nucleic Acids Res.* **40**, D370-D376 (2012)). A bound G protein is shown below the receptor (light grey). **b** Top view from the extracellular side, showing the ligand binding region of the receptors with labels for TM domains. **c** Bottom of the receptor from the cytosolic side, showing the G protein-binding region with TM domains labelled.

**Supplementary Fig. 2.**
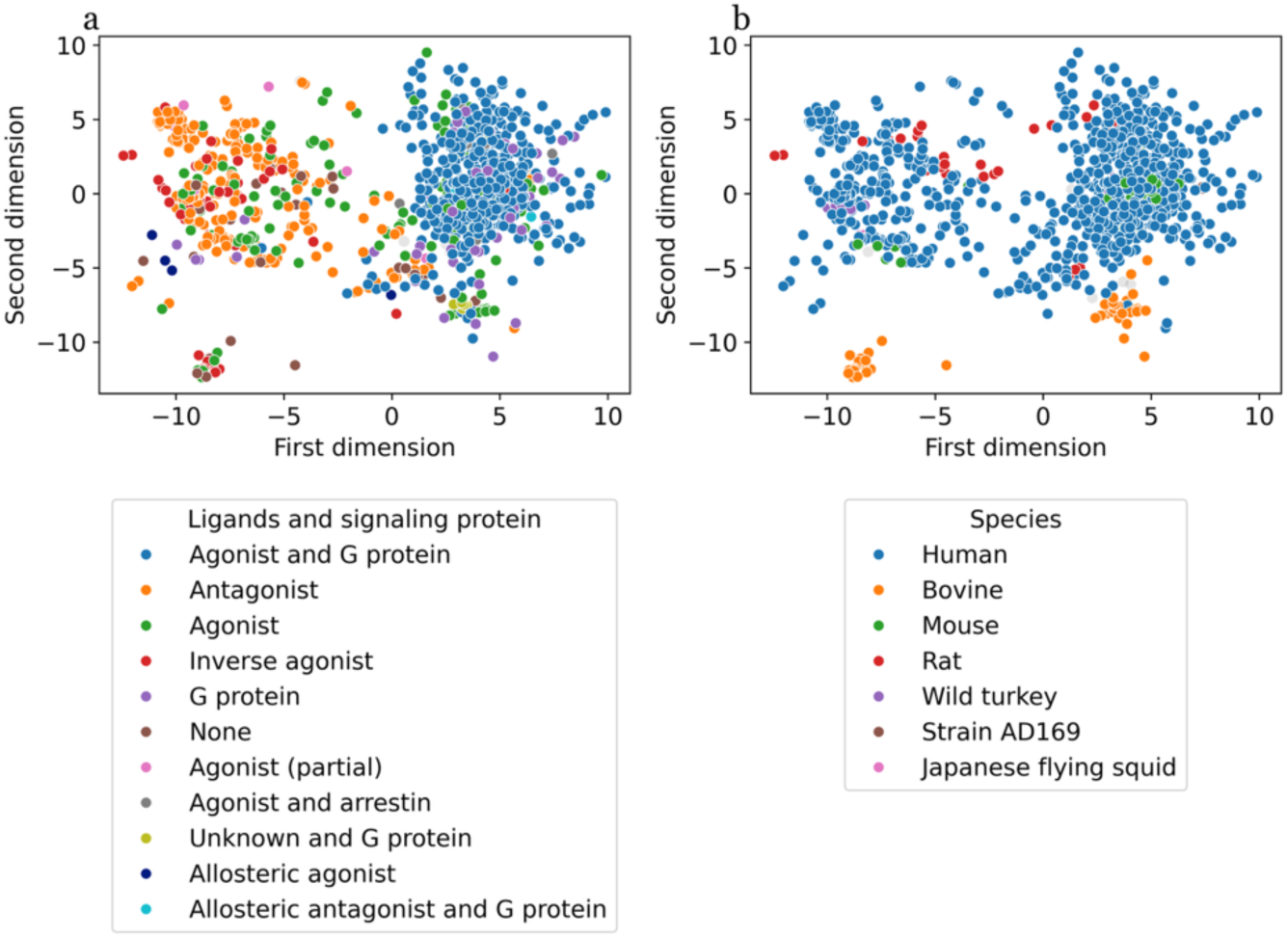
Scatter plots of the two first principal components in the PCA. **a** Illustration with colors reflecting whether and how the receptors are bound to small molecules. **b** Similar illustration but now with coloring based on the species. Only those categories with 5 or more datapoints are drawn here.

**Supplementary Fig. 3.**
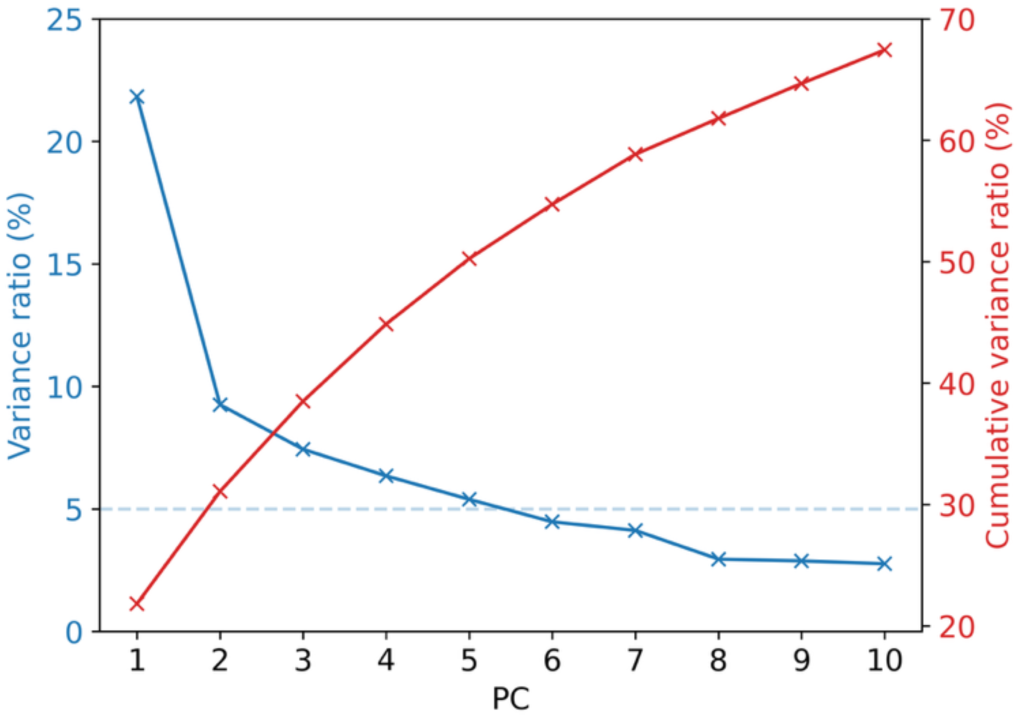
Variance ratio for the different principal components and the cumulative variance.

**Supplementary Fig. 4.**
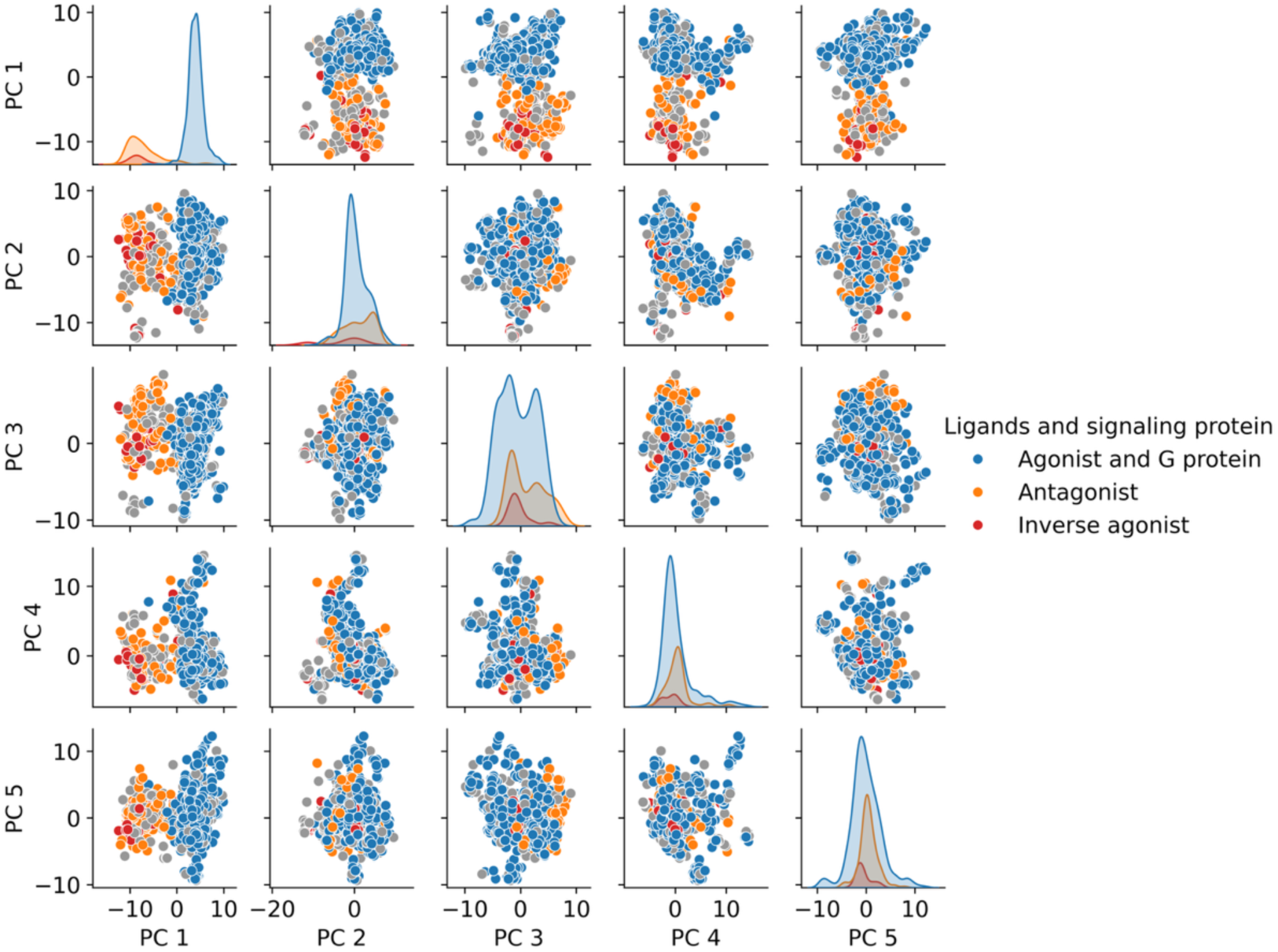
Pairwise scatter plots of projections along the first five principal components of the PCA. Colors indicate different bound states: antagonist bound (orange), agonist and G protein (blue), and other configurations (grey). Diagonal plots represent histograms of densities along each principal component. The first component (PC1) clearly separates the agonist and G protein-bound class from the antagonist-bound class.

**Supplementary Fig. 5.**
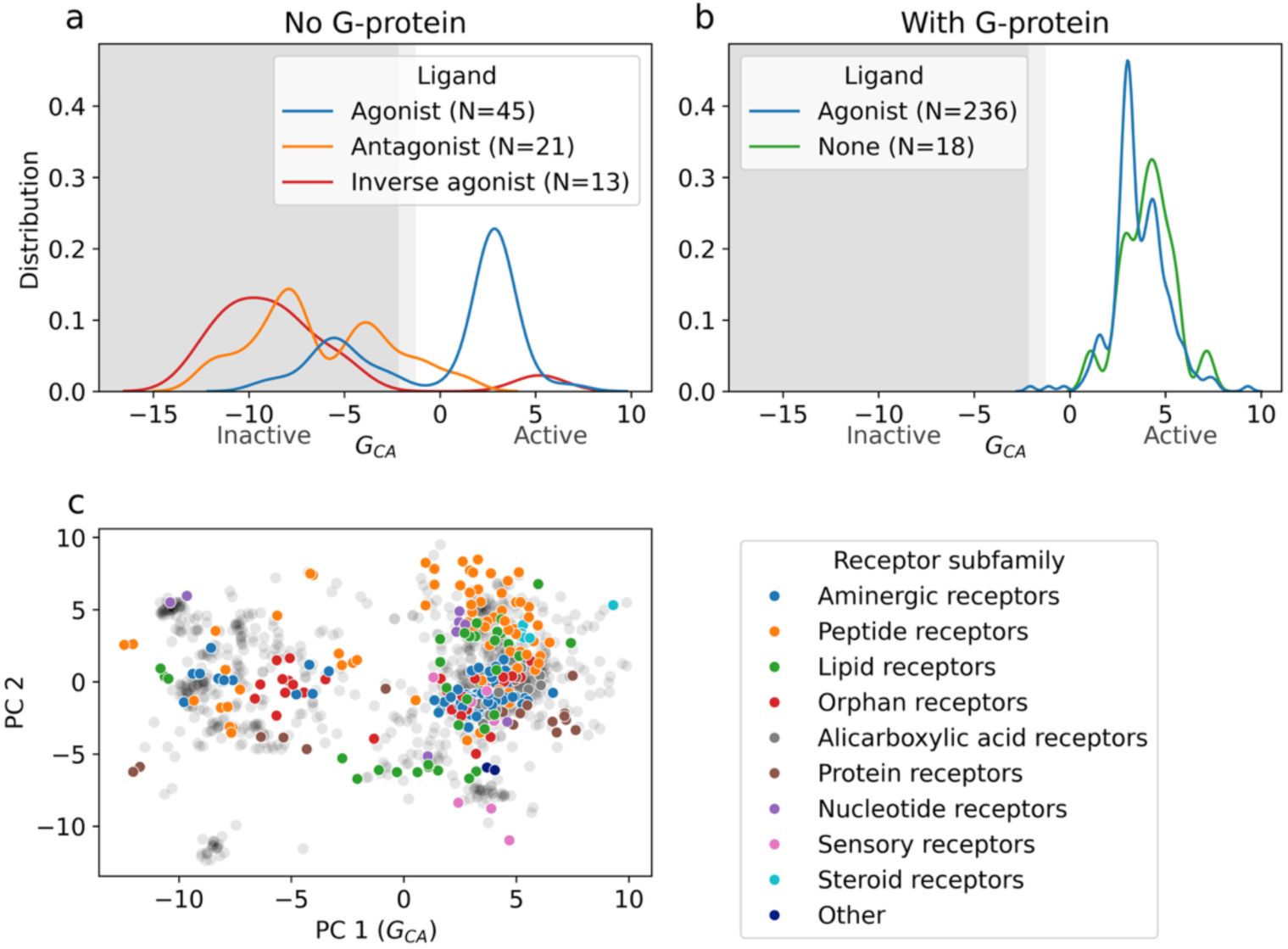
Results of the data not used for training. **a** Distributions along the index shown for new structures of ligand-bound receptors without G protein. **b** Distributions along the index shown for new structures of ligand-bound receptors with G protein. **c** Scatter plot of the data points of new structures shown over the first two principal components, colored by receptor subfamily. *N* stands for the number of structures analyzed.

**Supplementary Fig. 6.**
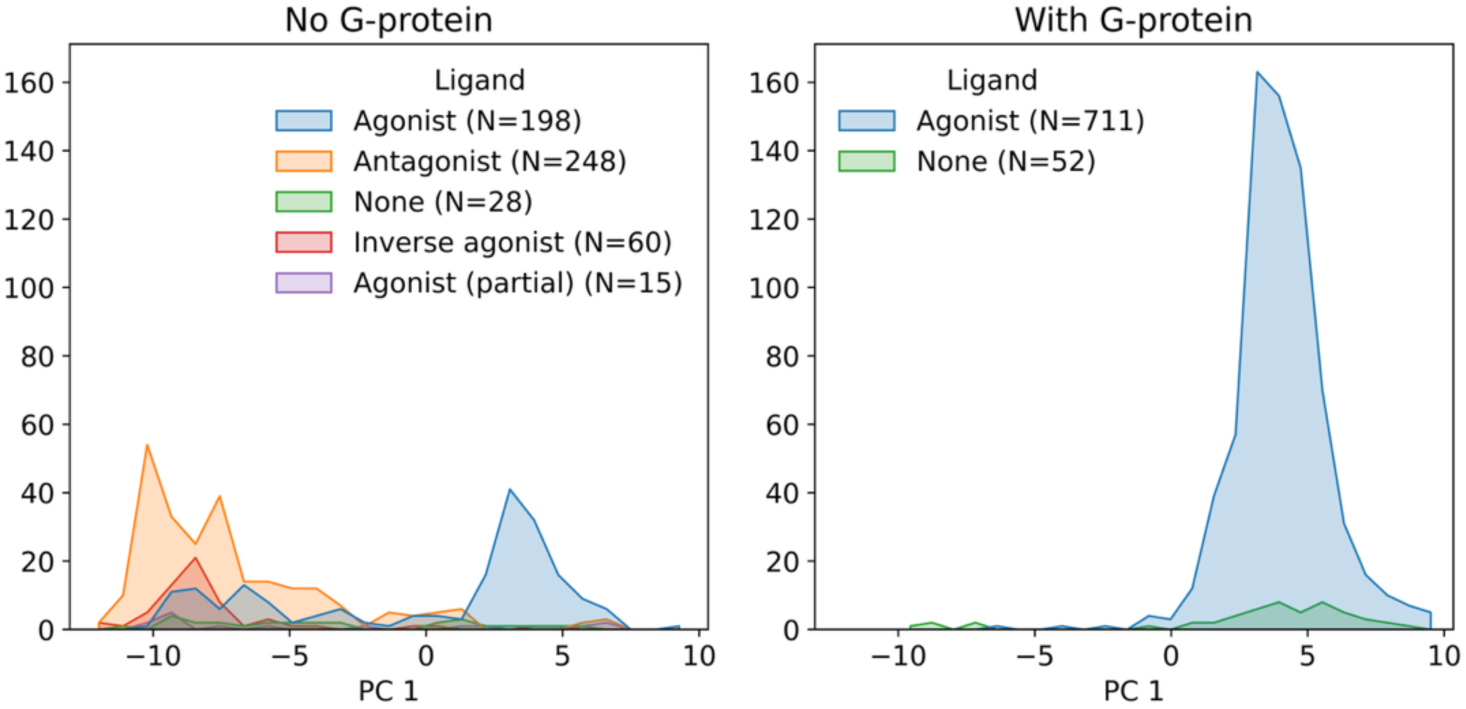
Influence of ligand in the distributions of receptor structures with and without a G-protein bound to the receptor, shown along the first principal component.

**Supplementary Table 1.** Number of structures expressed by receptor subfamily.

| SUBFAMILY | PEPTIDE<br>RECEPTORS | AMINERGIC<br>RECEPTORS | LIPID RECEPTORS | NUCLEOTIDE<br>RECEPTORS | ORPHAN<br>RECEPTORS | SENSORY<br>RECEPTORS | PROTEIN<br>RECEPTORS | ALICARBOXYLIC<br>ACID RECEPTORS | MELATONIN<br>RECEPTORS | OTHER | STEROID<br>RECEPTORS |
| --- | --- | --- | --- | --- | --- | --- | --- | --- | --- | --- | --- |
| TRAIN | 282 | 258 | 110 | 92 | 78 | 70 | 63 | 28 | 13 | 8 | 4 |
| TEST | 74 | 132 | 47 | 8 | 31 | 7 | 19 | 19 | 0 | 2 | 6 |
| TOTAL | 356 | 390 | 157 | 100 | 109 | 77 | 82 | 47 | 13 | 10 | 10 |

**Supplementary Table 2.**
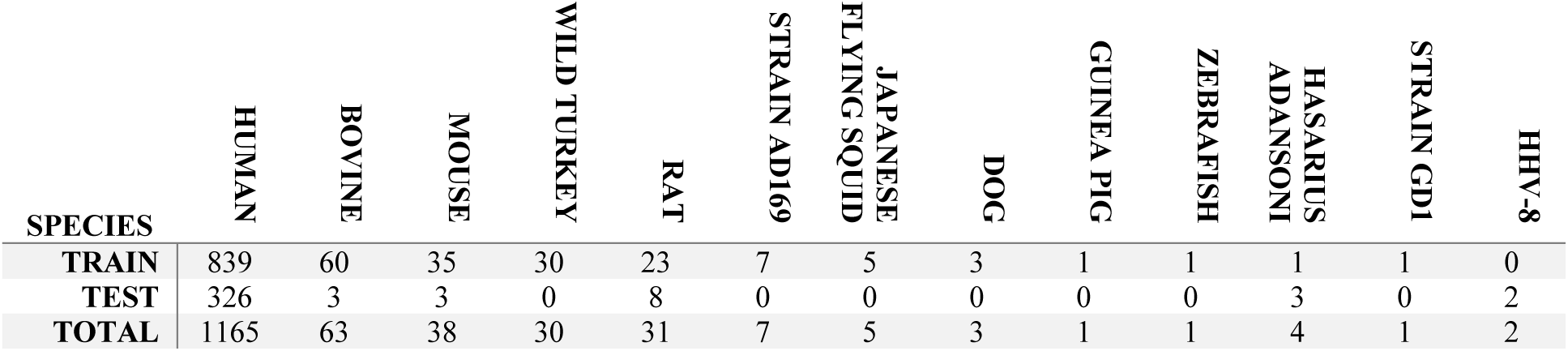
Number of structures expressed by species.

**Supplementary Table 3.** Number of structures expressed by which Ga is bound.

| Ga-type | None | G(i) | G(s) | G(q) | G(o) | G(t) | G-11 | G-13 | G(z) | G(olf) | G |
| --- | --- | --- | --- | --- | --- | --- | --- | --- | --- | --- | --- |
| TRAIN | 484 | 240 | 144 | 80 | 35 | 16 | 3 | 2 | 1 | 1 | 0 |
| TEST | 88 | 113 | 87 | 38 | 11 | 1 | 0 | 5 | 0 | 0 | 2 |
| TOTAL | 572 | 353 | 231 | 118 | 46 | 17 | 3 | 7 | 1 | 1 | 2 |

